# Rapamycin, an Immunosuppressant and mTORC1 Inhibitor, Triples the Antidepressant Response Rate of Ketamine at 2 Weeks Following Treatment: A double-blind, placebo-controlled, cross-over, randomized clinical trial

**DOI:** 10.1101/500959

**Authors:** Chadi G. Abdallah, Lynnette A. Averill, Ralitza Gueorguieva, Selin Goktas, Prerana Purohit, Mohini Ranganathan, Deepak Cyril D’Souza, Richard Formica, Steven M. Southwick, Ronald S. Duman, Gerard Sanacora, John H. Krystal

**Author notes:** Corresponding Author: Chadi G. Abdallah.

## Abstract

**BACKGROUND:** Ketamine exerts rapid and robust antidepressant effects thought to be mediated by activation of the mechanistic target of rapamycin complex 1 (mTORC1). To test this hypothesis, depressed patients were pretreated with rapamycin, an mTORC1 inhibitor, prior to receiving ketamine.

**METHODS:** Twenty-three patients suffering a major depressive episode were randomized to oral rapamycin (6 mg) or placebo, each was followed 2 hours later by ketamine 0.5 mg/kg in a double-blind cross-over design with treatment days separated by at least 2 weeks. Depression severity was assessed using Montgomery–Åsberg Depression Rating Scale (MADRS). Antidepressant response was defined as a MADRS improvement of 50% or more.

**RESULTS:** Over the two-week follow-up, we found a significant treatment by time interaction (F_(8,245)_ = 2.02, *p* = 0.04), reflecting prolonged antidepressant effects post rapamycin+ketamine treatment. At 2 weeks, we found a significantly higher response rate following rapamycin+ketamine (41%) compared to placebo+ketamine (13%, *p* = 0.04). However, rapamycin pretreatment did not alter the acute effects of ketamine.

**CONCLUSION:** Unexpectedly, pretreatment with rapamycin prolonged rather than blocked the acute antidepressant effects of ketamine. This observation raises questions about the role of mTORC1 in the antidepressant effects of ketamine, raises the possibility that rapamycin may extend the benefits of ketamine, and thereby potentially sheds light on mechanisms that limit the duration of ketamine effects. The supplementing of ketamine with rapamycin may be a treatment strategy for reducing the frequency of ketamine infusions during maintenance treatment.

## INTRODUCTION

Ketamine is an N-methyl-D-aspartate receptor (NMDAR) antagonist that exerts rapid and robust antidepressant effects (1,2). The antidepressant effects may emerge within hours of a single dose, but without additional ketamine doses, relapse typically occurs in 3-14 days (3-5). Ketamine and its metabolites are believed to exert antidepressant effects primarily by inducing a prefrontal glutamate neurotransmission surge leading to activation of synaptic α-amino-3-hydroxy-5-methyl-4-isoxazolepropionic acid glutamate receptors (AMPAR), which increases brain-derived neurotrophic factor (BDNF) levels, enhances stimulation of TrkB receptors, activates the mechanistic target of rapamycin complex 1 (mTORC1), and produces synaptogenesis (6-9). Several preclinical studies have shown that ketamine administration increases mTORC1 signaling (10-13), but there are non-replications of this finding (14). Most importantly, a single infusion of rapamycin into the medial prefrontal cortex (mPFC) prior to ketamine injection in rodents was reported to block the neuroplasticity and antidepressant-like effects of ketamine (10,15).

The current study tested the hypothesis that the antidepressant effects of ketamine are mediated by activation of mTORC1. This hypothesis was tested by evaluating whether the antidepressant effects of ketamine were blocked by pretreatment with the mTORC1 inhibitor, rapamycin. Following an experimental paradigm derived from animal research paradigms (10,15), we attempted to demonstrate in patients the observation that rapamycin blocks the antidepressant-like effects of ketamine (10,15).

In attempting to test the mTORC1 hypothesis of ketamine effects, we were aware of a major potential confound related to the anti-inflammatory effects of rapamycin. Neuroinflammation is increasingly implicated in the biology of depression, and anti-inflammatory effects of ketamine and other antidepressants may contribute to their antidepressant efficacy (16-20). Rapamycin is a powerful anti-inflammatory medication. Nonetheless these anti-inflammatory effects might augment those of ketamine, subsequently enhancing treatment efficacy. To detect possible synergistic effects, we followed patients for two weeks after each ketamine dose.

Using a randomized placebo-controlled cross-over design, rapamycin was administered as a single 6 mg dose prior to ketamine infusion. The rapamycin dose and timing were selected based on the drug pharmacokinetics to ensure, at the time of ketamine administration, blood concentration of 5 to 20 ng/mL, a level that exhibits potent immunosuppression (21). Consistent with the hypothesized mechanism of action of ketamine, we predicted that rapamycin would reduce the antidepressant effects of ketamine.

## METHODS AND MATERIALS

### Study Design

All study procedures were approved by an institution review board and all participants completed an informed consent process prior to enrollment (*Clinical-Trials.gov: NCT02487485*). A data and safety monitoring board (DSMB) oversaw the study protocol and monitored the study progress. The clinical trial included two study phases (I & II). In phase I, 3 participants received open-label rapamycin (a.k.a. sirolimus) followed 2 hr later by open-label ketamine. Participants remained on the research unit for at least 10 hr following the administration of rapamycin and were discharged upon clearance by the covering physician. The aim of phase I was to assess the safety and feasibility of the co-administration of rapamycin and ketamine.

In phase II, 23 participants were randomized to first receive either rapamycin or placebo, followed 2 hours later by open-label ketamine (See CONSORT Diagram in Supplements). Phase II was double-blind, placebo-controlled, cross-over design with at least 2 weeks between Infusion 1 (i.e., 1^st^ treatment day) and Infusion 2 (i.e., 2^nd^ treatment day). Depression severity no less than 20% of baseline was required prior to proceeding with Infusion 2. Participants who received placebo on Infusion 1 received rapamycin on Infusion 2, and vice-versa. Both study phases used a single-dose of 6mg rapamycin liquid form diluted in orange juice to maintain the blinding and ketamine 0.5mg/kg intravenously infused over 40 min. Ketamine administration and monitoring was comparable to previous studies (1,2,22).

Participants were assessed up to 2 weeks following each Infusion. Assessment measures included: 1) the Mini International Neuropsychiatric Interview (MINI) to determine the diagnosis, 2) Montgomery Åsberg Depression Rating Scale (MADRS) as primary outcome of depression severity, 3) Quick Inventory of Depressive Symptoms (QIDS) and Hamilton Anxiety Rating Scale (HAMA) as secondary measures of depression and anxiety severity, respectively, 4) Clinician Administered Dissociative States Scale (CADSS) and Positive and Negative Symptom Scale (PANSS), as safety measures of the psychotomimetic effects of ketamine, 5) rapamycin level immediately before starting ketamine and at the end of the ketamine infusion, and 6) high-sensitivity C-reactive protein (CRP) and erythrocyte sedimentation rate (ESR) prior to randomization to examine whether baseline inflammatory markers affect the antidepressant effects.

### Study Criteria

The study enrolled subjects between the age of 21 and 65 years. Participants were 1) diagnosed with current major depressive episode, 2) had a history of nonresponse to at least one adequate antidepressant trial, 3) were unmedicated or on a stable antidepressant or psychotherapy for at least 4 weeks, 4) had a MADRS ≥ 18 prior to randomization, 5) females were not pregnant or breastfeeding and were on a medically acceptable contraceptive method, 6) were able to read, write, and provide written informed consent, 7) did not have psychotic disorder or features, or manic or mixed episodes, 8) did not have an unstable medical condition, 9) did not require prohibited medications (see Table S1 in Supplements), 9) did not have urine drug screen positive for cannabis, phencyclidine, cocaine, or barbiturates, 10) had no substance dependence within 3 months, 11) had no sensitivity to rapamycin, ketamine, or heparin, and 12) had resting blood pressure higher than 85/55 and lower than 150/95 mmHg, and heart rate higher than 45/min and lower than 100/min.

### Statistics

Descriptive statistics (means, standard deviations and frequencies) were calculated prior to statistical analysis. Data distributions were checked using normal probability plots. The study a priori primary outcome was MADRS. Secondary outcomes included QIDS and HAMA, for depression and anxiety, respectively. Safety outcomes included CADSS and PANSS. Outcome variables were analyzed using mixed models with fixed effects of treatment (rapamycin vs. placebo), time (appropriate time points during Infusions 1 and 2), the interaction between treatment and time, and order (placebo first vs. rapamycin first). The best-fitting variance-covariance structure for each model was selected based on the Schwartz’ Bayesian Information criterion. Interactions between order and the other factors were checked for significance but not included in the final models for parsimony. Similarly, the effects of the variables CRP and ESR (log-transformed) were checked for significance but since non-significant were dropped from the final models. Post-hoc tests were used to interpret significant effects in the models: comparisons of treatment conditions by time-point for significant rapamycin by time interactions, and pairwise comparisons of time points for significant main effects of time. Least square means and standard errors by treatment and time, and by time were used for visualization of results. Response was defined as 50% improvement and its rate was compared between treatments using McNemar test. Effect size (Cohen’s d`) was calculated as the mean of the within-subject difference over its standard deviation. Correlation analyses explored the relationship between rapamycin level and improvement in depression severity.

The sample size was targeted based on feasibility within the 3-year funding available for this discovery phase project. Initially, we aimed to randomize 30 subjects in 3 years. However, we had a 1-year delay in starting randomization due to the addition of Phase 1 and the need for an investigational new drug exemption, both of which were requested by the institution review board. Thus, we were able to randomize a total of 23 patients in 2 years. Following randomization, one participant was excluded from the primary analysis due to receiving high dose hydrocortisone the night before randomization and the DSMB was informed accordingly. The decision to exclude the participant was made prior to compiling and unblinding the study data. However, for full transparency, a secondary analysis including this participant was conducted and reported in the Supplements. The results were found to be comparable to those of the primary analysis.

## RESULTS

### Participants

As detailed in the CONSORT Flow Diagram (see Supplements), 23 of the 57 assessed for eligibility were randomized and 20 participants were included in the analysis (2 subjects did not meet study criteria the morning of the first treatment day, and 1 subject received high dose hydrocortisone the night of the treatment day). The 20 participants were 8 men and 12 women, with mean (*±SEM*) age = 42.8 (*±2.8*) years, BMI = 27.2 (*±1.3*) kg/m^2^, CRP = 2.4 (*±0.8*) mg/L, ESR = 11.5 (*±2.3*) mm/hr, pre-infusion rapamycin = 26.5 (*±2.4*) ng/mL, and post-infusion rapamycin = 9.9 (*±1.0*) ng/mL.

### Treatment Effects on MADRS

MADRS was selected *a priori* as the primary outcome. There was a statistically significant interaction between treatment and time (F_(8,245)_ = 2.0, *p* = 0.04, Fig. 1A), with significant differences between rapamycin and placebo at day 3 (*p* = 0.04), and at day 5 (*p* = 0.02). There was also a significant main effect of time (F_(8,245)_ = 43.5, *p* < 0.0001), demonstrating significant decrease in MADRS scores from baseline, with the highest numerical mean difference achieved at 24 hr (17.5*±1.4*) and then gradually reduced until 2 weeks (8.5*±1.7*). However, the mean MADRS scores at 2 weeks remained significantly lower than baseline following both placebo (*Cohen’s d`* = 0.5; mean difference (*±SEM*) = 5.7 (*±2.5*), *t*_*(245)*_ = 2.3, *p* = 0.02) and rapamycin treatments (*Cohen’s d`* = 1.0; mean difference (*±SEM*) = 11.4 (*±2.4*), *t*_*(245)*_ = 4.7, *p* < 0.0001; Fig. 1A). There was no significant main effect of treatment (F_(8,245)_ = 1.4, *p* = 0.24) and the effects of the variables CRP and ESR were non-significant (*p* > 0.1). The response rates at 2 weeks were 13% following placebo and 41% following rapamycin treatment (*p* = 0.04; Fig. 1B).

**Figure 1.**
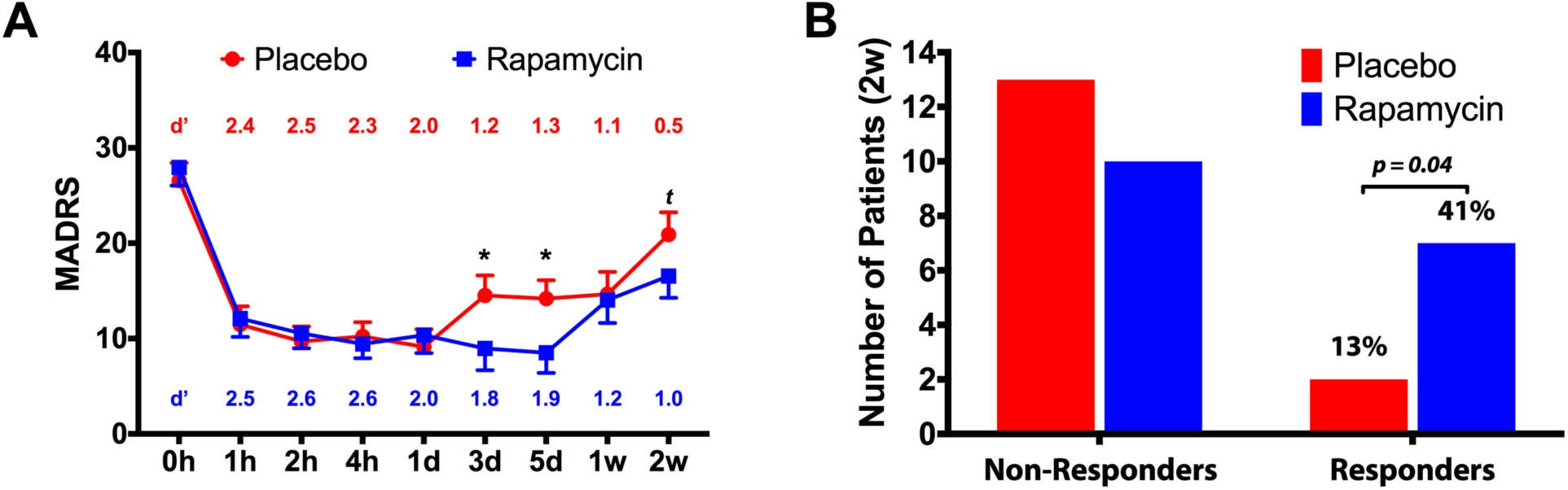
The Study Drug Effect on Montgomery Åsberg Depression Rating Scale (MADRS). **(A)** There is a significant main effect of time (F_(8,245)_ = 43.5, *p* < 0.0001), demonstrating significant decrease in MADRS scores from baseline. There is also significant interaction between treatment and time (F_(8,245)_ = 2.0, *p* = 0.04), with overall reduction in depression scores following treatment with rapamycin+ketamine (Rapamycin; blue line), compared to post placebo+ketamine (Placebo; red line). **(B)** Response rate was significantly higher following treatment with rapamycin+ketamine (Rapamycin; blue), compared to post placebo+ketamine (Placebo; red). *<u>NOTES</u>*: Response was defined as 50% improvement in MADRS from baseline; Error bars are standard errors of mean (*SEM*); d` = Cohen’s d` effect size compared to pretreatment MADRS scores; Comparison at each time point is marked with * for *p* ≤ 0.05 and *t* for *p* = 0.12;

### Treatment Effects on QIDS and HAMA

There was a significant main effect of time on QIDS (F_(8,236)_ = 7.1, *p* < 0.0001; Fig. S1), demonstrat-ing significant decrease in QIDS scores from baseline, with the highest numerical mean difference achieved at day 3 (5.2*±1.2*) and then gradually reduced until 2 weeks (2.1*±1.1*). The mean QIDS scores at 2 weeks remained significantly lower than baseline following rapamycin treatment (*Cohen’s d`* = 0.5; mean difference (*±SEM*) = 3.5 (*±1.5*), *t*_*(236)*_ = 2.4, *p* = 0.02), but not following placebo (*Cohen’s d`* = 0.1; mean difference (*±SEM*) = 0.7 (*±1.5*), *t*_*(236)*_ = 0.5, *p* = 0.64; Fig. S1). There was no significant main effect of treatment (F_(8,236)_ = 0.3, *p* = 0.57) or interaction between treatment and time (F_(8,236)_ = 0.5, *p* = 0.87).

There was a significant main effect of time on HAMA (F_(4,141)_ = 31.2, *p* < 0.0001), demonstrating significant decrease in HAMA scores from baseline, with the highest numerical mean difference achieved at 4 hr (9.9*±1.0*) and then gradually reduced until 2 weeks (3.0*±1.2*). The mean HAMA scores at 2 weeks was not significantly different compared to baseline following both placebo (*Cohen’s d`* = 0.4; mean difference (*±SEM*) = 3.0 (*±1.8*), *t*_*(141)*_ = 1.6, *p* = 0.11) and rapamy-cin treatments (*Cohen’s d`* = 0.4; mean difference (*±SEM*) = 3.1 (*±1.7*), *t*_*(141)*_ = 1.9, *p* = 0.07). There was no significant main effect of treatment (F_(1,141)_ = 0.2, *p* = 0.68) or interaction between treatment and time (F_(4,141)_ = 0.7, *p* = 0.63).

### Adverse Effects

There was a significant main effect of time on CADSS (F_(2,95)_ = 18.9, *p* < 0.0001), demonstrating significant increase in CADSS scores during infusion, which returned to baseline 2 hr post infusion (Fig. 2). There was no significant main effect of treatment (F_(2,95)_ = 0.2, *p* = 0.67) or interaction between treatment and time (F_(2,95)_ = 0.5, *p* = 0.60).

**Figure 2.**
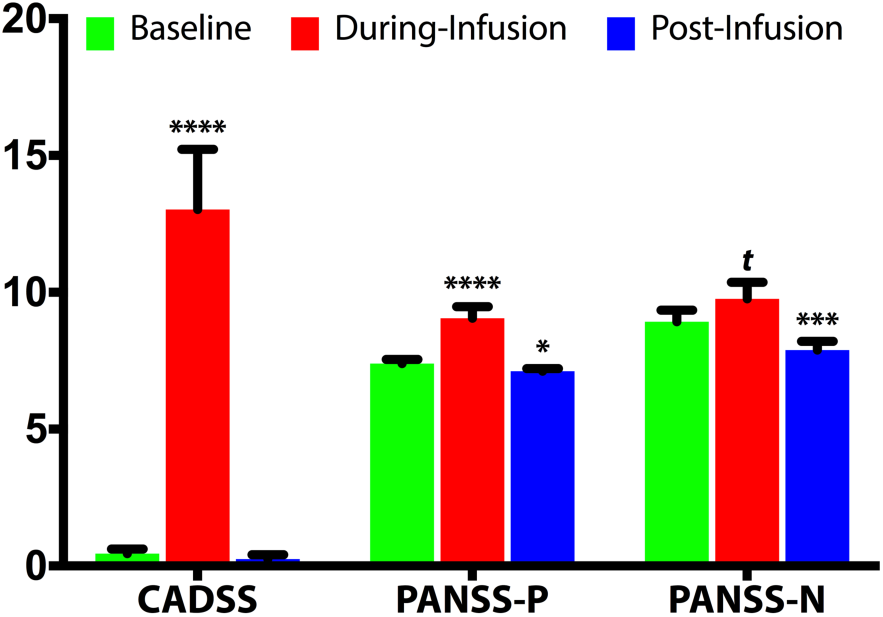
The Study Drug Effect on Clinician Administered Dissociative States Scale (CADSS), and Positive and Negative Symptom Scale (PANSS) Positive (PANSS-P) and Negative Symptoms (PANSS-N). Error bars are standard errors of mean (*SEM*); Comparisons to pretreatment scores are marked with **** for *p* ≤ 0.0001, *** for *p* ≤ 0.001, * for *p* ≤ 0.05 and ***t*** for *p* = 0.10.

There was a significant main effect of time on PANSS-positive (F_(2,82)_ = 11.3, *p* < 0.0001), demonstrating significant increase in PANSS-positive scores during infusion, with significant reduction 2 hr post infusion (Fig. 2). There was no significant main effect of treatment (F_(2,82)_ = 0.3, *p* = 0.57) or interaction between treatment and time (F_(2,82)_ = 1.9, *p* = 0.15). There was a significant main effect of time on PANSS-negative (F_(2,82)_ = 11.9, *p* < 0.0001), demonstrating significant reduction 2 hr post infusion (Fig. 2). There was no significant main effect of treatment (F_(1,82)_ = 0.2, *p* = 0.68) or interaction between treatment and time (F_(2,82)_ = 0.3, *p* = 0.73).

Participants in Phase 1 tolerated the combination treatment with no serious or unexpected adverse events. The study drug effects were clinically comparable to previous ketamine studies and there was no need for the extended monitoring of 10 hr. Therefore, we proceeded with the Phase 2 double-blind randomization and participants were discharged with transportation to home after medical clearance and completion of the last assessment on each treatment day. The adverse events during Phase 2 are reported in Table S2. There were no serious adverse events. New onset adverse events were mostly mild and transient. There were no reports of persistent adverse events. The most frequent adverse events were fatigue, headaches, nausea and pain. A total of 37 events were reported, 21 of which were reported by 4 participants.

## DISCUSSION

This study yielded two surprising, but potentially important, clinical observations. First, this study failed to validate the prediction from preclinical studies (10,15) in that rapamycin pretreatment did not reduce the acute antidepressant effects of ketamine. Second, rapamycin pretreatment tripled the response rate at 2 weeks, suggesting that this treatment approach may prolong the antidepressant effects of ketamine. This conclusion is supported by the statistically significant drug by treatment interaction effect on the primary outcome MADRS, showing overall larger reduction in depression scores following rapamycin pretreatment (Fig. 1A). Additionally, the Cohen’s d` effect size at 2 weeks post rapamycin was 1.0, compared to 0.5 following placebo pretreatment (Fig. 1A). Moreover, the reduction in QIDS scores (secondary outcome) at 2 weeks were significant following rapamycin, but not placebo pretreatment. This unanticipated finding is highly important considering the urgent need for treatment approaches to prolong the antidepressant effects of ketamine and other rapid-acting antidepressants. While infusions of ketamine 2 to 3 times per week have been shown to afford clinical benefit, less frequent administration is preferable to reduce the patient burden, adverse events, and drug abuse liability. Additionally, rapamycin pretreatment appears to have no effects on the anxiolytic or the psychotomimetic effects of ketamine. This suggests that the prolongation of the antidepressant effects of ketamine was not a consequence of changes in the subjective response to ketamine. Overall, rapamycin and ketamine were well tolerated with no serious adverse events.

Why are the antidepressant effects of ketamine transient and why are these effects prolonged by rapamycin? Briefly, one possibility suggested by this study is that ketamine treats depression without resolving underlying processes, such as neuroinflammation, that produce synaptic elimination and undermine antidepressant effects of ketamine. This hypothesis presumes that the expression of the antidepressant effects of ketamine depends upon sustaining the newly made synapses (6). The anti-inflammatory effects of rapamycin may protect these synapses and thereby extend the antidepressant effects of ketamine. A second hypothesis is a variant of the first in proposing that rapamycin may affect a homeostatic mechanism governing synaptic density. Both the detrimental effects of stress and the positive effects of ketamine upon synaptic density are transient, suggesting that these networks tend toward a stable level of synapses (23).

The synaptic model of depression is based on preclinical evidence that trauma and stress induce depression-like behavior along with a reduction in prefrontal synaptic density, and that both abnormalities are reversed by ketamine administration (10,15). Paralleling the preclinical findings, the evidence of prefrontal dysconnectivity in depression and normalization following ketamine treatment was also demonstrated in humans at the networks level (24-26). Yet, there are two critical observations of the synaptic model of stress that are less discussed and investigated. The first observation is that the stress-induced reduction in synaptic density is reversed to baseline within 2-4 weeks of stress cessation. The second observation is that the ketamine-induced increase in synaptic density also returns to baseline within 2 weeks of treatment (review in (6)). Together, this data raises the possibility of a subject-/region-specific synaptic density set point, highlighting the role of synaptic density stabilization in normal brain function as well as a target for treatment development. In the context of depression treatment, rather than administering ketamine repeatedly to increase synaptic density, ketamine might be administered to induce the reversal of synaptic density deficit and a complimentary drug might also be administered to ensure synaptic density stabilization at a new set point, and to prevent relapse.

The current study is unable to determine the mechanisms through which rapamycin prolonged the RAAD effects of ketamine. However, based on current understanding of the mechanisms of RAADs and rapamycin, a speculative model was developed to be tested in future studies. We hypothesize that the single rapamycin administration may have induced a transient synaptic density stabilization. Preclinical studies demonstrate that ketamine induction of synaptic density lasts for 7-10 days, which is paralleled by RAAD effects lasting 7-14 days (10,27). We hypothesize that rapamycin increases the stabilization of synaptic density stabilization for 3-5 days, which extended the RAAD effects of ketamine in a subgroup of treated patients. Notably, the half-life of rapamycin is approximately 60 hr and the administered rapamycin dose is expected to yield blood concentration that is approximately 1 ng/mL at days 3 to 5 (28). Our current working model is that the immunosuppressant actions of rapamycin induced synaptic density stabilization by reducing neuroinflammation. Disruption in the immune system has long been implicated in the pathology of depression, it remains to be seen in future mechanistic studies whether targeting this system may yield normalization of the synaptic density set point.

### Limitations and Strengths

As a first-in-humans study, the study sample was based on feasibility and funding availability rather than a priori knowledge of effect size. In addition, we were able to randomize patients for 2 years only, instead of 3 years due to the addition of Phase 1. Therefore, the lack of treatment by time interaction for QIDS may be the result of insufficient power to demonstrate a significant effect on this self-report measure of depression severity, which tends to have higher variability. Consistent with this possibility, the QIDS Cohen’s d` effect size at 2 weeks post rapamycin was 0.5, compared to only 0.1 following placebo treatment (Fig. S1). Based on our observation in Phase 1, we did not ask the participants to guess their treatment in Phase 2, as it was evident that the patients were unable to identify a rapamycin taste in the juice and the side effects were comparable to those seen in previous ketamine studies. Future studies may consider the benefit of adding an objective measure to determine the efficacy of the blinding.

A main strength of the study is the attempt to investigate an essential mechanistic pathway, that has been so far implicated in the pathology and treatment of depression based primarily on preclinical evidence. The results did not support the preclinical data, possibly due to the difference in route of drug delivery (i.e., preclinical studies used intracerebral infusion of rapamycin into the mPFC). Other strengths of the study include a double-blind randomized placebo-controlled design. The cross-over also afforded a within subject comparison, further highlighting the contrast between the two treatments. Finally, the fact that the immune system is involved in both depression pathology as well as in resilience and depression recovery (29) creates a major challenge in the field, emphasizing the need to target a “sweet spot” that will oppose the negative effects of inflammation while avoiding the inhibition of its neuroregulatory function (29). Therefore, an essential strength of the study is the use of combined therapy, instead of monotherapy or add-on approaches that were used in the past (18). If successfully developed as one drug administration every 7-14 days, combined therapy will overcome many of the short-comings of anti-inflammatory monotherapy/add-on approaches, including: 1) lower burden and less adverse effects compared to daily use of drugs, and 2) the effect appears to be independent of pretreatment exaggerate inflammatory state (i.e., no CRP or ESR effects), which appears to be necessary for successful monotherapy/add-on approaches (30).

## CONCLUSION

The administration of a single dose of rapamycin, reaching blood levels known to induce potent immunosuppression, does not inhibit the RAAD effects of ketamine. Intriguingly, the immunosuppressant rapamycin prolonged the antidepressant effects of ketamine and tripled the response rate at 2 weeks following treatment. To date, preclinical and clinical studies based on the synaptic model of depression have largely focused on the transient alteration in synaptic density. Future studies providing greater insight into the mechanisms of synaptic density stabilization and approaches to alter the putative synaptic density set point may provide novel target for drug development and could ultimately lead to depression cure rather than treatment.

## ACKNOWLEDGMENTS

The authors would like to thank the individuals who participated in this study and the members of the data and safety monitoring board (DSMB) who have overseen the study protocol and progress. The authors also thank the Emerge Research Program staff and the nurses and lab staff of the Biostudies Unit for the invaluable expertise in conducting this trial. Funding and research support were provided by Pfeiffer Foundation, NIMH (K23MH101498), and the VA National Center for PTSD, the CT Department of Mental Health and Addiction Services and Yale-New Haven Health. The content is solely the responsibility of the authors and does not necessarily represent the official views of the sponsors, the Department of Veterans Affairs, NIH, or the U.S. Government.

## SUPPLEMENTAL INFORMATION

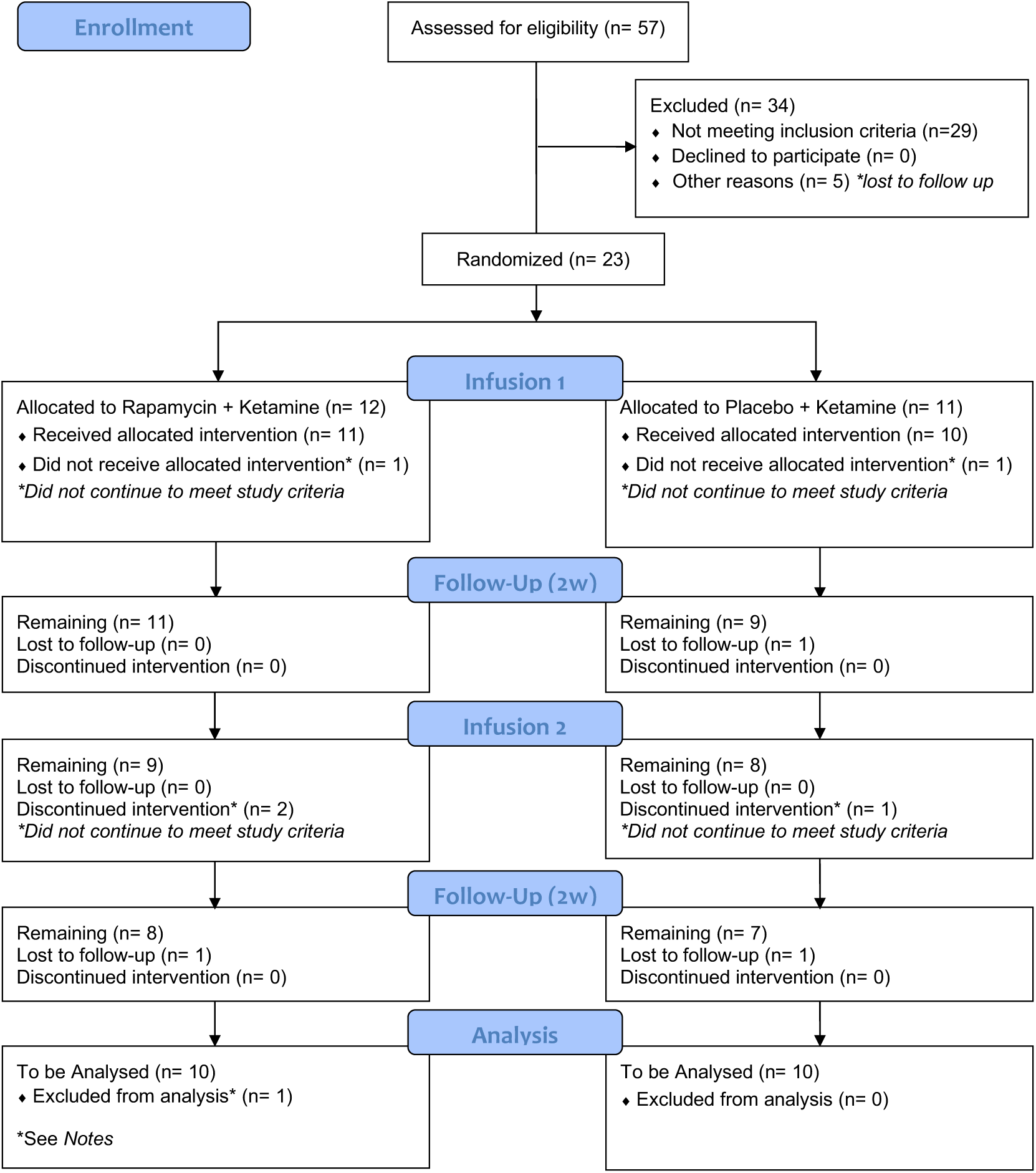

## Additional Analyses

Here we repeated the primary analysis after including the subject who was excluded because of taking a large dose of hydrocortisone the night of treatment day, investigating the effects of the study drug on the primary outcome, Montgomery Åsberg Depression Rating Scale (MADRS). The results were similar to those found in the primary analysis. There was a statistically significant interaction between treatment and time (F_(8,261)_ = 2.3, *p* = 0.02). There was also a significant main effect of time (F_(8,261)_ = 49.0, *p* < 0.0001). There was no significant main effect of treatment (F_(8,261)_ = 1.4, *p* = 0.25).

**Figure S1.**
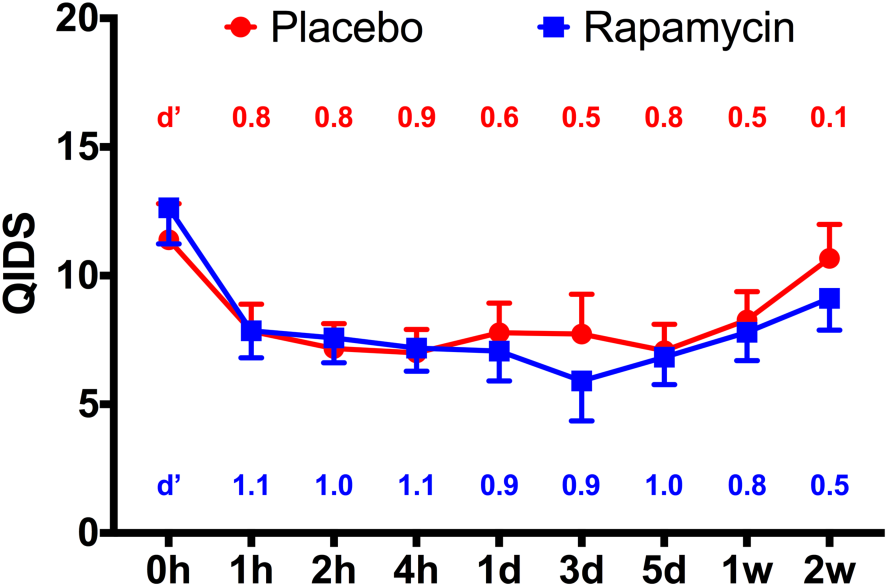
The Study Drug Effect on Quick Inventory of Depressive Symptoms (QIDS). There is a significant main effect of time on QIDS (F_(8,236)_ = 7.1, *p* < 0.0001), demonstrating significant decrease in QIDS scores from baseline following treatment with either rapamycin+ketamine (Rapamycin; blue line) or placebo+ketamine (Placebo; red line). There is no statistically significant interaction between treatment and time (F_(8,236)_ = 0.5, *p* = 0.87). Error bars are standard errors of mean (*SEM*); d` = Cohen’s d` effect size compared to pretreatment QIDS scores;

**Table S1.**
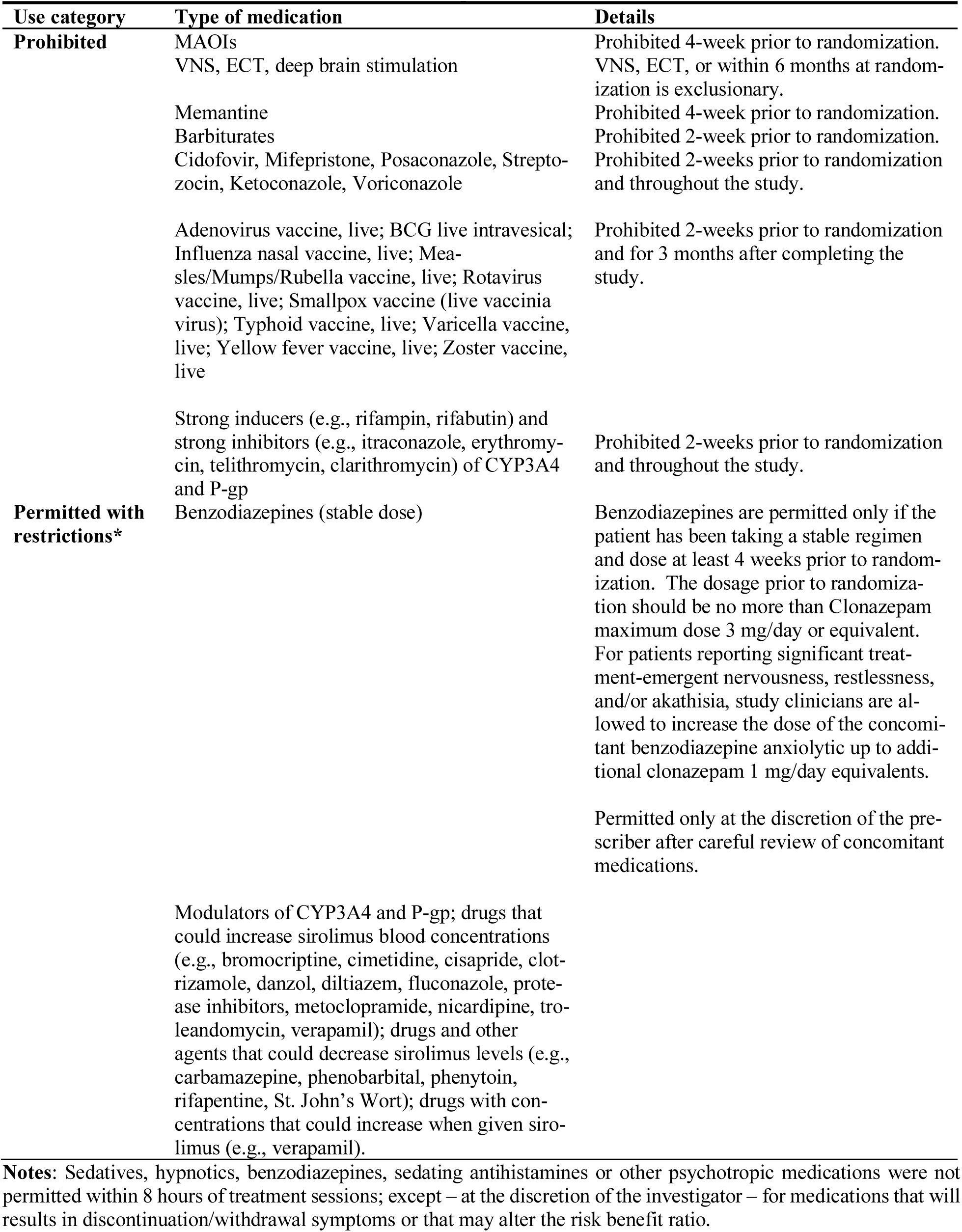
Concomitant Treatments that are prohibited

**Table S2.**
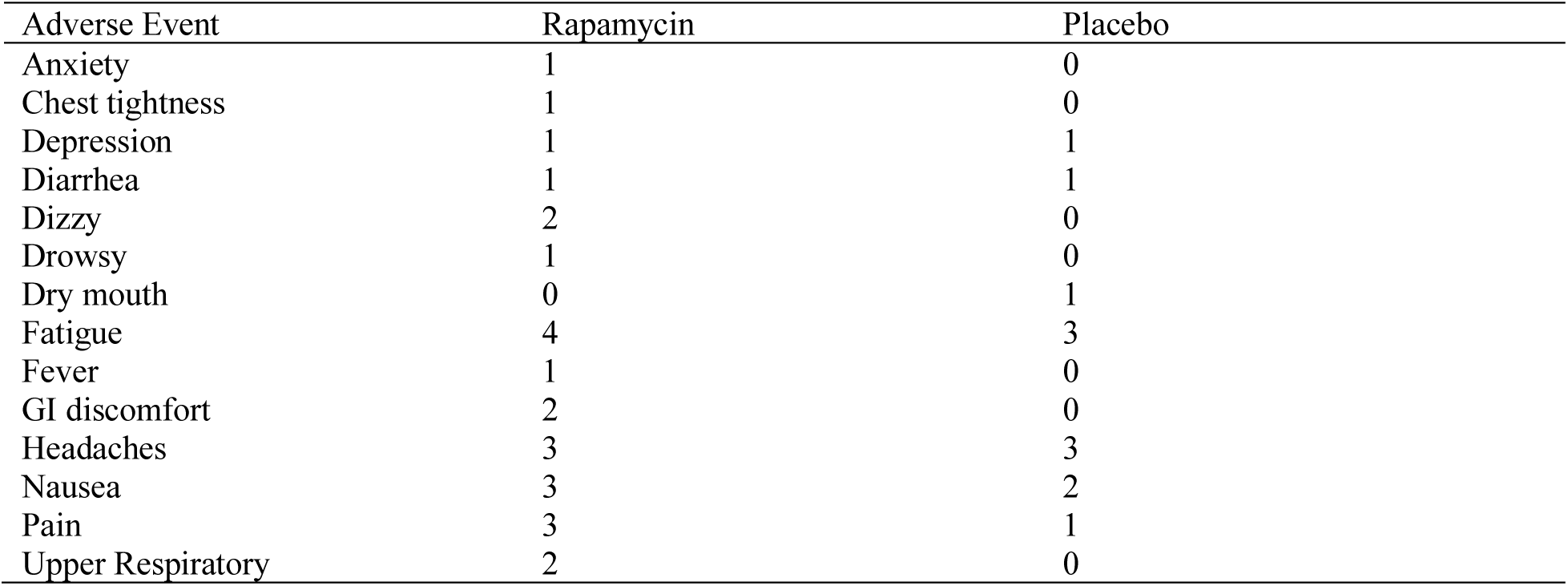
Averse Events Profile

| Adverse Event | Rapamycin | Placebo |
| --- | --- | --- |
| Anxiety | 1 | 0 |
| Chest tightness | 1 | 0 |
| Depression | 1 | 1 |
| Diarrhea | 1 | 1 |
| Dizzy | 2 | 0 |
| Drowsy | 1 | 0 |
| Dry mouth | 0 | 1 |
| Fatigue | 4 | 3 |
| Fever | 1 | 0 |
| GI discomfort | 2 | 0 |
| Headaches | 3 | 3 |
| Nausea | 3 | 2 |
| Pain | 3 | 1 |
| Upper Respiratory | 2 | 0 |

## REFERENCES

1. Murrough JW, Iosifescu DV, Chang LC, Al Jurdi RK, Green CE, Perez AM, et al. (2013): Antidepressant efficacy of ketamine in treatment-resistant major depression: a two-site randomized controlled trial. Am J Psychiatry. 170:1134–1142.

2. Zarate CA, Jr., Singh JB, Carlson PJ, Brutsche NE, Ameli R, Luckenbaugh DA, et al. (2006): A randomized trial of an N-methyl-D-aspartate antagonist in treatment-resistant major depression. Arch Gen Psychiatry. 63:856–864.

3. Aan Het Rot M, Zarate CA, Jr., Charney DS, Mathew SJ (2012): Ketamine for depression: where do we go from here? Biol Psychiatry. 72:537–547.

4. Abdallah CG, Averill LA, Krystal JH (2015): Ketamine as a promising prototype for a new generation of rapid-acting antidepressants. Ann N Y Acad Sci. 1344:66–77.

5. Romeo B, Choucha W, Fossati P, Rotge JY (2015): Meta-analysis of short- and mid-term efficacy of ketamine in unipolar and bipolar depression. Psychiatry Res. 230:682–688.

6. Abdallah CG, Sanacora G, Duman RS, Krystal JH (2018): The neurobiology of depression, ketamine and rapid-acting antidepressants: Is it glutamate inhibition or activation? Pharmacol Ther.

7. Gould TD, Zarate CA, Jr., Thompson SM (2018): Molecular Pharmacology and Neurobiology of Rapid-Acting Antidepressants. Annu Rev Pharmacol Toxicol.

8. Murrough JW, Abdallah CG, Mathew SJ (2017): Targeting glutamate signalling in depression: progress and prospects. Nat Rev Drug Discov. 16:472–486.

9. Ignacio ZM, Reus GZ, Arent CO, Abelaira HM, Pitcher MR, Quevedo J (2016): New perspectives on the involvement of mTOR in depression as well as in the action of antidepressant drugs. British journal of clinical pharmacology. 82:1280–1290.

10. Li N, Lee B, Liu RJ, Banasr M, Dwyer JM, Iwata M, et al. (2010): mTOR-dependent synapse formation underlies the rapid antidepressant effects of NMDA antagonists. Science. 329:959–964.

11. Zhou W, Wang N, Yang C, Li XM, Zhou ZQ, Yang JJ (2014): Ketamine-induced antidepressant effects are associated with AMPA receptors-mediated upregulation of mTOR and BDNF in rat hippocampus and prefrontal cortex. Eur Psychiatry. 29:419–423.

12. Yang C, Hu YM, Zhou ZQ, Zhang GF, Yang JJ (2013): Acute administration of ketamine in rats increases hippocampal BDNF and mTOR levels during forced swimming test. Ups J Med Sci. 118:3–8.

13. Harraz MM, Tyagi R, Cortes P, Snyder SH (2016): Antidepressant action of ketamine via mTOR is mediated by inhibition of nitrergic Rheb degradation. Mol Psychiatry. 21:313–319.

14. Popp S, Behl B, Joshi JJ, Lanz TA, Spedding M, Schenker E, et al. (2016): In search of the mechanisms of ketamine’s antidepressant effects: How robust is the evidence behind the mTor activation hypothesis. F1000Research. 5.

15. Li N, Liu RJ, Dwyer JM, Banasr M, Lee B, Son H, et al. (2011): Glutamate N-methyl-D-aspartate receptor antagonists rapidly reverse behavioral and synaptic deficits caused by chronic stress exposure. Biol Psychiatry. 69:754–761.

16. Haroon E, Miller AH, Sanacora G (2017): Inflammation, Glutamate, and Glia: A Trio of Trouble in Mood Disorders. Neuropsychopharmacology. 42:193–215.

17. Miller AH (2013): Conceptual confluence: the kynurenine pathway as a common target for ketamine and the convergence of the inflammation and glutamate hypotheses of depression. Neuropsychopharmacology. 38:1607–1608.

18. Kohler O, Benros ME, Nordentoft M, Farkouh ME, Iyengar RL, Mors O, et al. (2014): Effect of anti-inflammatory treatment on depression, depressive symptoms, and adverse effects: a systematic review and meta-analysis of randomized clinical trials. JAMA psychiatry. 71:1381–1391.

19. Kadriu B, Gold PW, Luckenbaugh DA, Lener MS, Ballard ED, Niciu MJ, et al. (2018): Acute ketamine administration corrects abnormal inflammatory bone markers in major depressive disorder. Mol Psychiatry. 23:1626–1631.

20. Walker AK, Budac DP, Bisulco S, Lee AW, Smith RA, Beenders B, et al. (2013): NMDA receptor blockade by ketamine abrogates lipopolysaccharide-induced depressive-like behavior in C57BL/6J mice. Neuropsychopharmacology. 38:1609–1616.

21. Mahalati K, Kahan BD (2001): Clinical pharmacokinetics of sirolimus. Clinical pharmacokinetics. 40:573–585.

22. Berman RM, Cappiello A, Anand A, Oren DA, Heninger GR, Charney DS, et al. (2000): Antidepressant effects of ketamine in depressed patients. Biol Psychiatry. 47:351–354.

23. Abdallah CG, Averill LA, Akiki TJ, Raza M, Averill CL, Gomaa H, et al. (2018): The Neurobiology and Pharmacotherapy of Posttraumatic Stress Disorder. Annu Rev Pharmacol Toxicol.

24. Abdallah CG, Averill CL, Salas R, Averill LA, Baldwin PR, Krystal JH, et al. (2017): Prefrontal Connectivity and Glutamate Transmission: Relevance to Depression Pathophysiology and Ketamine Treatment. Biol Psychiatry Cogn Neurosci Neuroimaging. 2:566–574.

25. Abdallah CG, Averill LA, Collins KA, Geha P, Schwartz J, Averill C, et al. (2017): Ketamine Treatment and Global Brain Connectivity in Major Depression. Neuropsychopharmacology. 42:1210–1219.

26. Abdallah CG, Dutta A, Averill CL, McKie S, Akiki TJ, Averill LA, et al. (2018): Ketamine, but Not the NMDAR Antagonist Lanicemine, Increases Prefrontal Global Connectivity in Depressed Patients. Chronic Stress. 2:2470547018796102.

27. Liu RJ, Fuchikami M, Dwyer JM, Lepack AE, Duman RS, Aghajanian GK (2013): GSK-3 inhibition potentiates the synaptogenic and antidepressant-like effects of subthreshold doses of ketamine. Neuropsychopharmacology. 38:2268–2277.

28. Brattstrom C, Sawe J, Jansson B, Lonnebo A, Nordin J, Zimmerman JJ, et al. (2000): Pharmacokinetics and safety of single oral doses of sirolimus (rapamycin) in healthy male volunteers. Therapeutic drug monitoring. 22:537–544.

29. Miller AH, Haroon E, Felger JC (2017): Therapeutic Implications of Brain-Immune Interactions: Treatment in Translation. Neuropsychopharmacology. 42:334–359.

30. Raison CL, Rutherford RE, Woolwine BJ, Shuo C, Schettler P, Drake DF, et al. (2013): A randomized controlled trial of the tumor necrosis factor antagonist infliximab for treatment-resistant depression: the role of baseline inflammatory biomarkers. JAMA psychiatry. 70:31–41.

